# Identification of a distal RXFP1 gene enhancer with differential activity in fibrotic lung fibroblasts involving AP-1

**DOI:** 10.1101/2021.07.23.453556

**Authors:** Ting-Yun Chen, Xiaoyun Li, Gillian C. Goobie, Ching-Hsia Hung, Kyle Hamilton, Harinath Bahudhanapati, Jiangning Tan, Daniel J. Kass, Yingze Zhang

## Abstract

Relaxin/insulin-like family peptide receptor 1 (RXFP1) mediates relaxin’s antifibrotic effects and has reduced expression in the lung and skin of patients with fibrotic interstitial lung disease (fILD) including idiopathic pulmonary fibrosis (IPF) and systemic sclerosis (SSc). This may explain the failure of relaxin-based anti-fibrotic treatments in SSc, but the regulatory mechanisms controlling *RXFP1* expression remain largely unknown. This study aimed to identify regulatory elements of *RXFP1* that may function differentially in fibrotic fibroblasts.

We identified and evaluated a distal regulatory region of *RXFP1* in lung fibroblasts using a luciferase reporter system. Using serial deletions, an enhancer upregulating pGL3-promoter activity was localized to the distal region between -584 to -242bp from the distal transcription start site (TSS). This enhancer exhibited reduced activity in IPF and SSc lung fibroblasts. Bioinformatic analysis identified two clusters of activator protein 1 (AP-1) transcription factor binding sites within the enhancer. Site-directed mutagenesis of the binding sites confirmed that only one cluster reduced activity (−358 to -353 relative to distal TSS). Co-expression of FOS in lung fibroblasts further increased enhancer activity. *In vitro* complex formation with a labeled probe spanning the functional AP-1 site using nuclear proteins isolated from lung fibroblasts confirmed a specific DNA/protein complex formation. Application of antibodies against JUN and FOS resulted in the complex alteration, while antibodies to JUNB and FOSL1 did not. Analysis of AP-1 binding in 5 pairs of control and IPF lung fibroblasts detected positive binding more frequently in control fibroblasts. Expression of *JUN* and *FOS* was reduced and correlated positively with *RXFP1* expression in IPF lungs.

In conclusion, we identified a distal enhancer of *RXFP1* with differential activity in fibrotic lung fibroblasts involving AP-1 transcription factors. Our study provides insight into *RXFP1* downregulation in fILD and may support efforts to reevaluate relaxin-based therapeutics alongside upregulation of *RXFP1* transcription.

## Introduction

Pulmonary fibrosis is a hallmark of fibrotic interstitial lung diseases (fILD). Although the pathogenesis of fILD is not fully understood [1], fibroblast activation in the lungs of patients with fILD results in aberrant extracellular matrix (ECM) collagen accumulation [2]. Idiopathic pulmonary fibrosis (IPF) and systemic sclerosis (SSc) are two of the most common types of fILD. IPF is a chronic and progressive disease associated with high morbidity and mortality [1, 2]. In patients with SSc, fILD is the disease manifestation associated with the highest mortality [3]. Despite the increasing global burden of fILDs [4, 5], our understanding of the mechanisms underlying the development and progression of fibrosis and our ability to target these pathogenic pathways is lacking.

Relaxin is a heterodimeric peptide hormone that regulates collagen metabolism and ECM turnover [6]. Relaxin was considered to be a potent anti-fibrotic agent [7-11], but clinical studies have failed to show beneficial anti-fibrotic effects in patients with SSc [12]. Relaxin mediates its cellular effects through its receptor, Relaxin/insulin-like family peptide receptor 1 (RXFP1) [13]. It plays an important homeostatic role in tissue remodeling, for example through collagen relaxation of pelvic ligaments during parturition [14]. In fibrotic diseases, the relaxin/RXFP1 axis is dysregulated [14]. *RXFP1*-null mice develop early onset peribronchiolar and perivascular fibrosis, with *relaxin* knock out mice developing early pulmonary and systemic organ fibrosis [15, 16]. *RXFP1* expression is downregulated in whole lung tissue and lung fibroblasts from patients with fILD, including IPF and SSc [17-20]. *In vitro* studies of fibroblasts isolated from IPF and SSc lungs demonstrates minimal responsiveness to relaxin treatment in reducing extracellular matrix accumulation, but restoration of *RXFP1* expression restores the anti-fibrotic effects in these cells [17]. However, transcriptional regulation of *RXFP1* in fibroblasts is poorly understood. Characterization of *RXFP1* regulation will provide insight to therapeutic targets for restoring relaxin’s anti-fibrotic effects in patients with fILD [14].

Activator protein 1 (AP-1) belongs to the superfamily of basic leucine zipper DNA-binding transcription factors. It exists as a dimer mainly consisting of two subfamilies: Fos and Jun subunits [21]. AP-1 targets the TPA response element (TRE, also known as the AP-1 site) that regulates gene expression in response to physiologic and pathologic functions [22]. This includes the transcriptional upregulation of genes important for tissue remodeling [23]. AP-1 also plays a central role in enhancer repertoires selection in fibroblasts, which are critical for tissue differentiation during development [24]. There is limited research to date investigating the role of AP-1 superfamily transcription factor regulation of *RXFP1*.

In this study, we sought to characterize the regulatory regions of the *RXFP1* gene and to identify transcriptional elements important in its regulation. Through fine mapping of these regions, we identified a novel distal enhancer containing specific binding motifs for AP-1. We further demonstrated direct binding of AP-1 to the *RXFP1* regulatory elements using *in vitro* models. Our study provides insight to the transcriptional regulation of *RXFP1* in lung fibroblasts, which may have future implications for relaxin-based therapeutics.

## Methods

### Cell Culture

The study was approved and was determined to be “non-human” study by the Institutional Review Board at the University of Pittsburgh (STUDY18100070). Donor lungs were obtained from the CORE (Center for Organ Recovery and Education). IPF and SSc explanted lungs were recovered from patients who underwent lung transplantation at the University of Pittsburgh Medical Center. Lung fibroblast lines derived from these lungs were maintained in Dulbecco’s modified Eagle’s medium (DMEM) with 10% fetal calf serum and 50 μg/mL penicillin-streptomycin (Thermo Fisher Scientific Inc.) at 37°C and 5% CO_2_.

### Plasmids and Cloning

Polymerase chain reaction (PCR) products were gel purified using Qiagen QlAquick gel purification columns (Qiagen) according to the manufacturer’s instructions. The PCR products were cloned using promoter-less pGL3-basic (pB) vector or pGL3-promoter (pP) vector containing a SV40 promoter (Promega Corporation) and Gibson Assembly (New England BioLabs). The relative location and size of *RXFP1* DNA in each luciferase reporter plasmid are listed in **Supplemental Table 1**.

### Dual Luciferase Assay

Fibroblasts were seeded at 5,000 cells/well in 24 well cell culture plates and cultured overnight prior to transfection with either the pGL3-*RXFP1* reporter plasmids alone (0.4μg/well) or co-transfection with a transcription factor expression plasmid (0.3μg pGL3-*RXFP1* reporter and 0.1μg expression plasmid per well). A Renilla luciferase vector (pGL4.74 [hRluc/TK]) was used as a control (0.001μg/well, Promega) for transfection efficiency. Plasmids were transfected into primary lung fibroblasts using Lipofectamine 2000 according to the manufacturers’ instruction. At 40 hours post-transfection, the cells were washed with PBS, lysed in 1 × passive lysis buffer and analyzed using the Dual-Luciferase Reporter Assay System (Promega Corporation) and a SpectraMax L Microplate Reader (Molecular Devices, LLC.). Relative expression levels of pGL-*RXFP1* reporters were normalized against pB or pP vector luciferase activity.

### Prediction of putative promoter and TATA element

DNA sequences upstream of both distal and proximal transcriptional start sites (TSS) were used to identify putative promoter and TATA elements. The Neural Network Promoter Prediction method (http://www.fruitfly.org/seq_tools/promoter.html) was used with a minimum promoter score of 0.85 [25]. The location of each identified element was determined based on the corresponding TSS.

### Site-Directed Mutagenesis

Site-directed mutagenesis of the AP-1 binding sites in the distal enhancer reporter plasmids were performed using the Q5® Site-Directed Mutagenesis Kit (New England BioLabs). Primer pairs 5’-CCATAATGTGgggCTATACTAAATTTCATCTTC-3’ and 5’-CTAAATCCACTTAGAAAAAACAATC-3’; 5’-AGCATGCATGgggCACAGATTGTTC-3’ and 5’-AAATGTAGCCAAACCCAG-3’ were used for binding site 1 and site 2, respectively.

### Nuclear Protein Extraction

Nuclear proteins were prepared using fibroblasts at 80-90% confluency and the Nuclear Extract Kit (Active Motif), according to the manufacturer’s protocol.

### Electrophoretic Mobility Shift Assays (EMSA) and Supershift Analysis

A 37-base pair (bp) double-stranded oligonucleotide containing the AP-1-binding motif (underlined) of the *RXFP1* enhancer (5’-TACATTTAGCATGCA<u>TGACTC</u>ACAGATTGTTCTAGA-3’) was used as a probe for EMSA. The probe was biotin labeled at the 3’ end using a Pierce™ Biotin 3’ End DNA Labeling Kit (Thermo Fisher Scientific Inc.). Seven micrograms of nuclear protein were added to a binding reaction mixture containing 2µl 10X binding buffer, 1µl poly (dl·dC), 1µl 50% glycerol, 1µl 1% NP-40 with 20fmol biotin-labeled probe and incubated at room temperature for 20 minutes. After incubation, 5µL of 5X loading buffer was added to each binding reaction mixture and 20μL was loaded into each well for polyacrylamide gel electrophoresis. Nylon membrane was used for gel transfer. DNA was crosslinked using UV light and detected using the LightShift(tm) Chemiluminescent EMSA Kit (Thermo Fisher Scientific Inc.). For competition binding reaction, unlabeled wildtype, described above, or mutated AP-1 binding site probe (5’-TACATTTAGCATGCATGgggCACAGATTGTTCTAGA-3’) was added in 50-fold excess to the reaction mixture. Supershift assays were performed by incubating monoclonal antibody (Ab) to specific AP-1 transcription factors c-Jun (60a8, Cell Signaling), c-Fos (9F6), FOSL1(Fra-1, D-3), and JUNB (C-11) from Santa Cruz Biotechnology with nuclear proteins for 10 minutes on ice and 10 minutes at room temperature prior to the binding reaction described above. Rabbit IgG (Cell Signaling) was used as a negative control.

### Chromatin Immunoprecipitation (ChIP) Assay

ChIP assay was performed as described [26, 27]. Briefly, lung fibroblasts were grown on 100-mm tissue culture dishes to 90% confluence. Cells were cross-linked with 1% formaldehyde for 10 minutes and harvested for fragmentation using sonication. The chromatin fragments were immunoprecipitated with 3μg of the indicated antibodies for c-JUN (Cell signaling) and rabbit normal IgG (Cell signaling). The precipitated fragments were washed five times and analyzed by PCR using a primer pair (F: 5’-AAACACTGGACTGGGTTTGG-3’ and R: 5’-GGAAAGTAGGCCCCTTGAGA-3’) spanning the putative AP-1 binding site 2 on the *RXFP1* enhancer. ChIP assay was performed using rabbit IgG as a negative control. Densitometry analysis of the PCR amplification was performed using ImageJ [28] Positive pulldown of bound DNA sequences was determined using the IgG as a control.

### RXFP1 and AP-1 gene expression

Lung tissue-specific expressions of *RXFP1, JUN*, and *FOS* genes (where *JUN and FOS* are both AP-1 transcription factors) were obtained from the publicly available Lung Genomics Research Consortium (LGRC) gene expression dataset (GEO accession GSE47460; http://www.lung-genomics.org/) [29]. *FOS* gene expression was analyzed using microarray and was available for 108 controls and 160 IPF patients. *JUN* gene expression was only available from the RNA sequencing (RNAseq) dataset for 22 controls and 22 IPF patients. The expression levels on RNAseq were shown in Fragments Per Kilobase of transcript per Million mapped reads (fPKM) and were square root transformed for normality prior to analysis.

### Statistical Analysis

All data were expressed as the mean ± SD. Student’s *t-*test was used for two-way comparisons. Gene expression levels of control and IPF groups were compared using the Mann-Whitney U test. Correlation of *FOS* and *JUN* gene expression levels with *RXFP1* expression levels was analyzed using linear regression modeling as described [30]. All analyses were performed in Prism GraphPad version 7.05 and a *p* value < 0.05 significance threshold was used.

## Results

### Identification of a functional promoter associated with distinct RXFP1 transcripts

*RXFP1* is located on chromosome 4 (hg38, chr4:158,521,714-158,653,367) and historically was thought to be solely comprised of a 132kb (kilobase pair) genomic sequence (designated as the “Short” form of *RXFP1*). Subsequently the GENCODE project reported one extended *RXFP1* transcript with 206.4kb additional sequences (hg38, chr4:158,315,311-158,652,212) upstream of the previously reported transcript as shown in the University of California Santa Cruz (UCSC) genome browser (https://genome.ucsc.edu/). This is designated as the “Long” form of *RXFP1* (https://www.gencodegenes.org/). As shown in **Figure 1A**, there are multiple splicing variants associated with Short *RXFP1* [31], while only one transcript is associated with the Long form.

**Figure 1.**
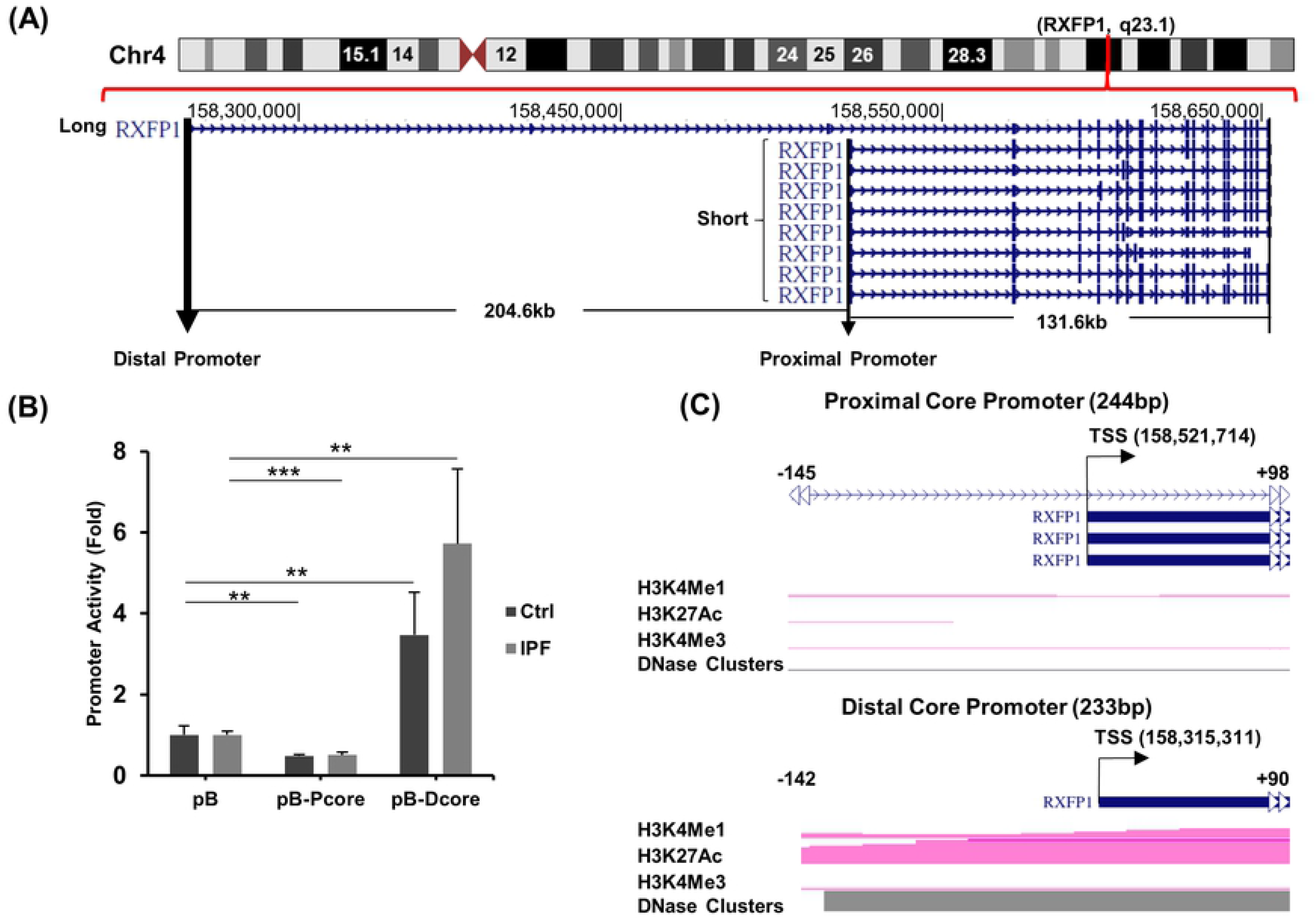
The genomic structure of *RXFP1* and core promoter activity. **(A)** Location of the Long and Short forms of *RXFP1* with identification of their mapped transcripts on chromosome 4 is shown. Genomic locations were determined based on the human genome build hg38 (https://genome.ucsc.edu/). The putative proximal and distal promoters are shown. **(B)** Proximal and distal core (P-Core and D-core, respectively) promoter activity analysis using a Luciferase promoter reporting system is shown. Transfections were performed in quadruplicates using Lipofectamine 2000 (Thermo-Fisher). Both control (Ctrl) and idiopathic pulmonary fibrosis (IPF) lung fibroblasts were used. The promoter activity (fold) was calculated using pGL3basic (pB) as a control. Student T-test (two tailed) were used for the pairwise comparison of luciferase activity with p-values reported as: *<0.05, **<0.01, ***<0.001. Mean and standard deviation are shown for each plasmid. **(C)** Chromatin characteristics associated with active transcriptional regulation including H3K4Me1, H3K27Ac, H3K4Me3 and DNAse sensitivity clusters for the P-Core and D-Core region using the Encyclopedia of DNA Elements (ENCODE) histone ChIP data (UCSC genome browser) are shown. The nucleotide location relative to each of the transcription start site (TSS) are labelled. *RXFP1*, Relaxin/insulin-like family peptide receptor 1; UCSC, University of California, Santa Cruz.

To determine whether a functional promoter is associated with each of the two forms of *RXFP1*, we analyzed the core promoter regions of each transcript using a pGL3 luciferase reporter system and primary lung fibroblasts isolated from donor lungs, as controls, and IPF lungs. A 233bp DNA element spanning -142 to +90 of the distal TSS (hg38, chr4:158,315,311) for the Long form (distal promoter), and a 194bp fragment covering -145 to +48 of the proximal TSS (hg38, chr4:158,521,714) for the Short form (proximal promoter) were tested (hereafter, all sequence locations are numbered relative to its corresponding TSS). As shown in **Figure 1B**, the distal promoter showed increased activity compared with pB vector, a promoter-less vector for testing promoter activity of targeted sequences, in both control and IPF lung fibroblasts (p=0.004 and 0.002, respectively). In contrast, the reporter activities for the proximal promoter in both control and IPF fibroblasts were reduced compared with the pB vector.

We further analyzed the two promoter regions for chromatin characteristics associated with active transcriptional regulation including H3K4Me1, H3K27Ac, H3K4Me3 and DNAse sensitivity clusters using the Encyclopedia of DNA Elements (ENCODE) histone ChIP data tracts in the UCSC genome browser (**Figure 1C**). Consistent with the reporter assay, only the distal promoter region was associated with positive transcriptional regulation signals, indicating that this was the only functional core promoter for the *RXFP1* gene in lung fibroblasts.

### Differential distal promoter activities between control and fibrotic lung fibroblasts

Given the lack of promoter activity in the core proximal regulatory region, we extended our search for potential regulatory elements to both the proximal (PE) and distal (DE) regulatory regions. We analyzed the likelihood of a functional promoter by identifying a TATA box in a 3.1kb region (−2158bp to +971bp) and a 1.4kb region (−1202bp to +161bp) within the DE and PE regions, respectively. These regions possess potential regulatory functions based on the UCSC genome browser. Consistent with the lack of proximal promoter activity, there was no TATA box within 200bp upstream of the proximal TSS. However, a potential site was identified in the proximal region at -1095 to -1114. For the distal region, a TATA box was identified at -16 to +3 in addition to another site between -1946bp to -1927bp (**Figure 2A**).

**Figure 2.**
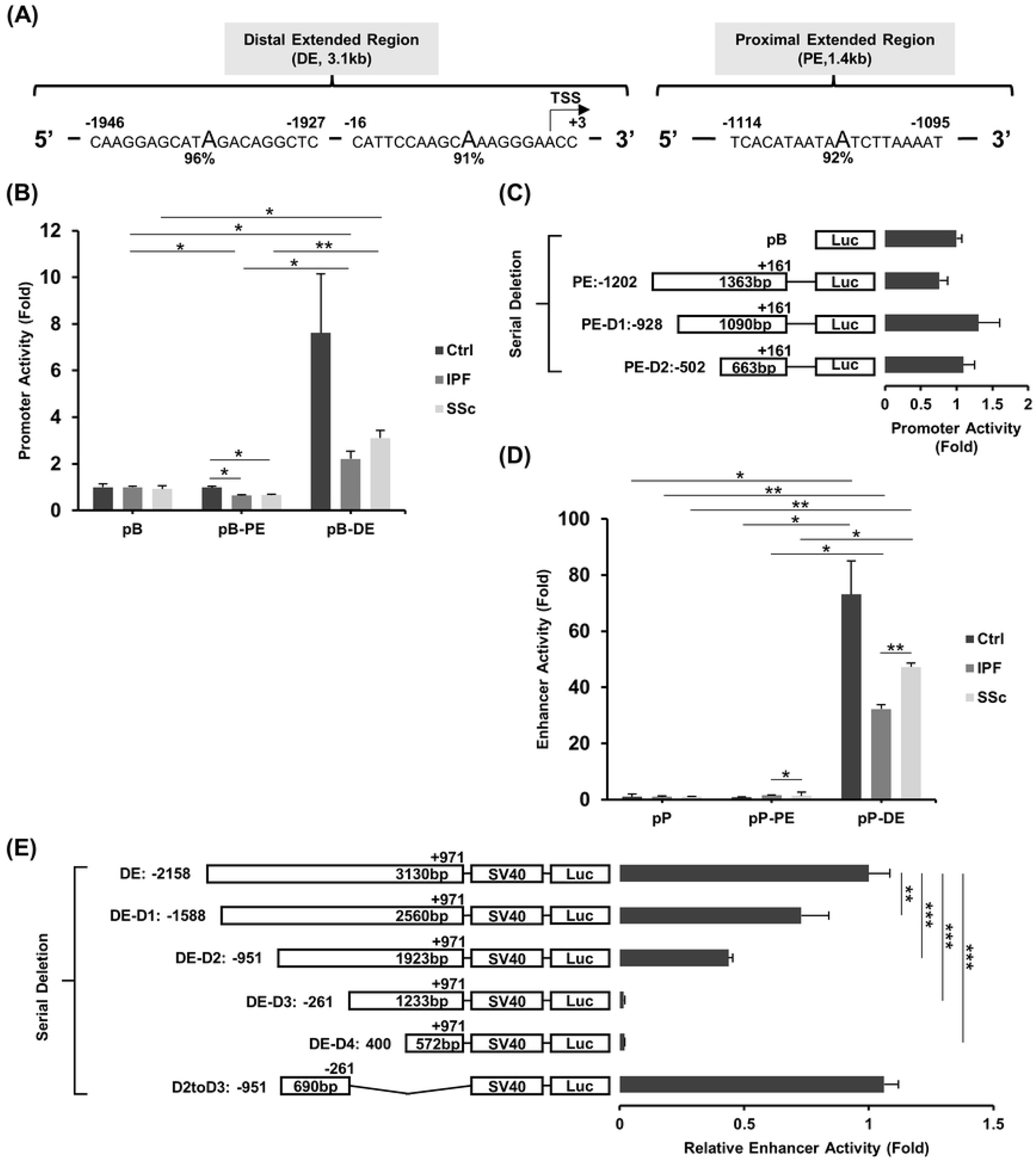
The promoter and enhancer activity in the extended proximal and distal regulatory regions. **(A)** Locations of TATA elements identified in the extended 3.1kb distal (DE) and 1.4kb proximal (PE) regulatory regions is shown using the Neural Network Promoter Prediction method (http://www.fruitfly.org/seq_tools/promoter.html). Relative nucleotide locations to the TSS are labelled. **(B)** Promoter activity of the extended proximal (pB-PE) and distal (pB-DE) regulatory regions with pGL3 promoter reporter system (transfections performed in duplicates) using control (Ctrl), idiopathic pulmonary fibrosis (IPF) and systemic sclerosis (SSc) lung fibroblasts is shown. Fold changes are relative to the pGL3basic (PB) vector. **(C)** Deletional analyses of the extended proximal regulatory region (PE-D1, proximal deletion 1; PE-D2, proximal deletion 2) using control fibroblasts (transfections performed in triplicates) are shown. Fold changes are relative to the pGL3basic (pB) vector. **(D)** Enhancer activity of the extended proximal and distal regulatory regions with pGL3 enhancer reporter system (transfections performed in duplicates) using control, IPF and SSc lung fibroblasts are shown. Fold changes are relative to the pGL3promoter (pP) vector. **(E)** Serial deletion of the extended distal regulatory region is shown. The deletion plasmids are sequentially labeled as DE-D1 to DE-D4, and the plasmid with the sequence between DE-D2 and DE-D3 retained and a deletion of the retained sequence in DE-D3 is labeled as DE-D2toD3. The beginning and ending locations relative to the TSS and fragment size for each clone are labelled. Relative enhancer activities are calculated by obtain the fold change using pP as a control for each reporter plasmid and subsequently using this to calculate the fold change between each plasmid to the original 3.1kb DE plasmid. For **(B)**-**(E)**, student’s T-test (two tailed) was used for pairwise comparisons of luciferase activity. P-values are reported as: *<0.05, **<0.01, ***<0.001. Mean and standard deviation are shown for each plasmid.

These extended regions were further characterized in lung fibroblasts from control, IPF, and SSc patients for promoter activity using pB. Similar to the proximal core promoter, there was no increased activity for the PE in any fibroblast lines compared to the pB vector (**Figure 2B**). The 3.1kb DE retained its promoter activity and higher promoter activity was observed in control (7.6 ± 2.5 fold) compared to either IPF (2.2 ± 0.3 fold) or SSc (3.1 ± 0.3 fold) fibroblasts, although neither reached statistical significance (p=0.096 and 0.129, respectively).

We performed serial deletions of the 1.4kb PE to rule out any repressor element interfering with promoter activity. Deleting 274 bp or 700 bp upstream sequences did not result in any significant promoter activity increase compared to the pB vector (**Figure 2C**), further supporting that only the distal regulatory region has promoter function.

### Localization of an enhancer region upstream of the distal promoter with differential activities in control and fibrotic fibroblasts

Since the extended distal region retained promoter activities among control and fibrotic fibroblasts, we tested whether the extended distal region was associated with enhancer function using pGL3promoter, which contains a SV40 promoter and used for testing enhancer activity of targeted sequences. Using control fibroblasts, we consistently observed greater than 50-fold enhancer activity in the distal region while there was no activity for the proximal extended region compared to the pP vector (**Figure 2D**). The distal enhancer activity was significantly reduced in IPF (32.3 ± 1.56 fold, p=0.041) and SSc (47.3 ± 1.3 fold, p=0.042) fibroblasts compared to controls (73.1 ± 11.9 fold). Deletion of 570bp (DE-D1), 637bp (DE-D2) in the 5’ sequences of the 3.1kb extended distal region retained 73 ± 11% and 44 ± 2% enhancer activity (p=0.008 and <0.001, respectively) (**Figure 2E**). Deletion of an additional 690bp (DE-D3) and 1360bp (DE-D4) completely abolished the enhancer activity (p<0.001 for both), suggesting that the enhancer is localized to a 690bp region between -951 to -261 (designated as the distal *RXFP1* enhancer). This was confirmed with the additional deletion of 1233bp 3’ sequences (DE-D2toD3) of the D2 clone that fully restored the 3.1kb enhancer activity (106 ± 6%).

### Fine mapping of the distal enhancer region

The distal enhancer partially overlaps with a region of dense transcription factor binding sites (TFBS, https://genome.ucsc.edu/)(**Figure** 3A). Therefore, we constructed a 608bp (−675 to -68) clone based on the TF binding cluster and designated it as pP-TFBS. Direct comparison of the distal *RXFP1* enhancer (pP-D2toD3) and the TFBS element showed similar enhancer activities in control and SSc lung fibroblasts (**Figure 3B**).

**Figure 3.**
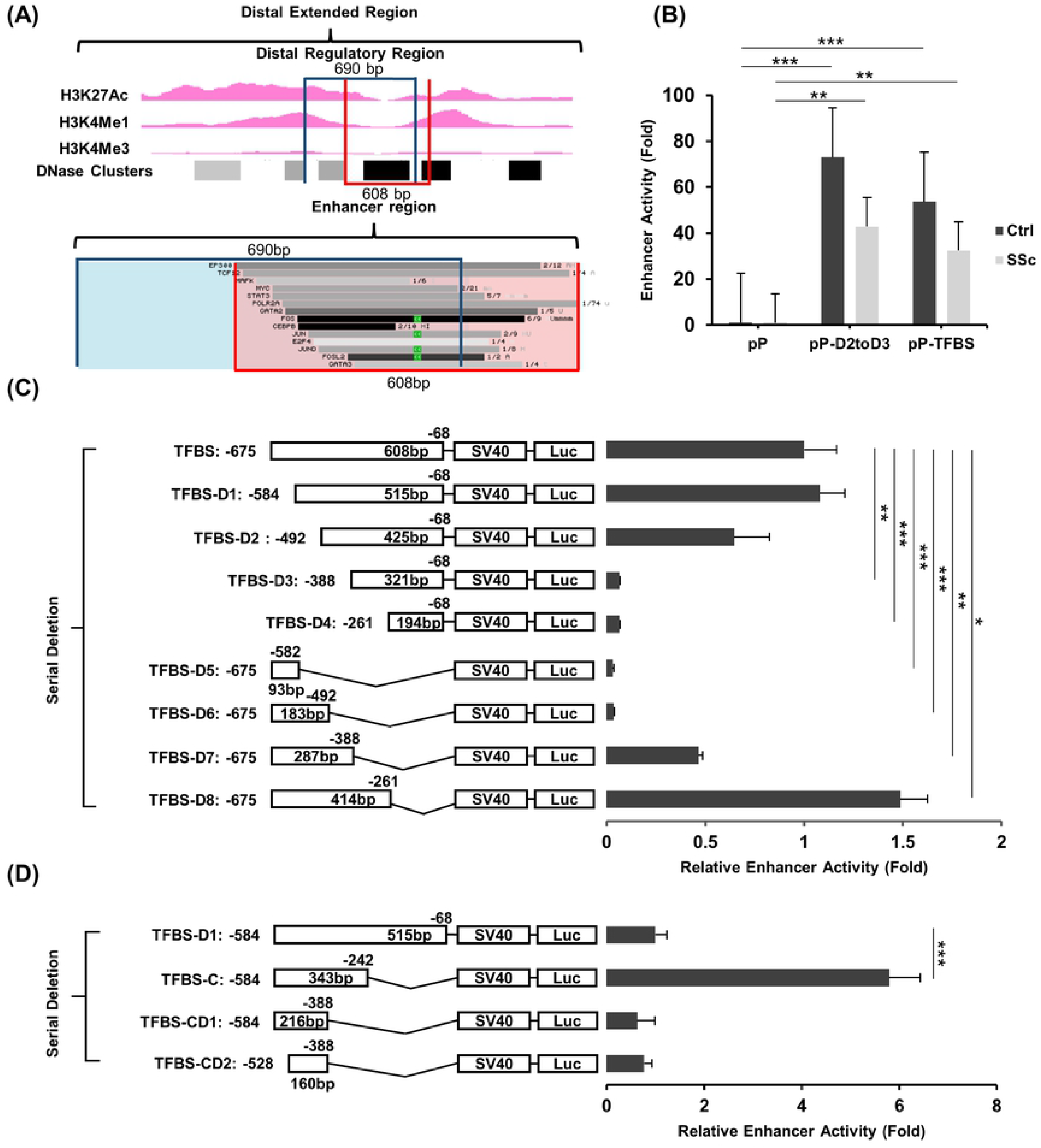
Fine mapping of the distal enhancer. **(A)** Chromatin characteristics including H3K4Me1, H3K27Ac, H3K4Me3 and DNAse sensitivity clusters for the region flanking the 690bp distal enhancer and the transcription factor binding sites (TFBSs) using UCSC genome browser are shown. **(B)** Enhancer activity of the 690bp (pP-D2toD3) and 608bp (pP-TFBS) distal enhancer (transfections performed in triplicates) using control and SSc lung fibroblasts with the pGL3promoter (pP) as a control are shown. **(C)** and **(D)** Serial deletion of the 608bp (pP-TFBS) and 515bp (TFBS-D1) distal enhancer are shown. In **(C)** the deletion plasmids are sequentially labeled as TFBS-D1 to D8 and in **(D)**, they are labelled as TFBS-C and TFBS-CD1 to CD2. Transfections were performed in triplicates. Relative enhancer activities are calculated by obtaining the fold change using pP as a control for each deletion plasmid and subsequently using this to calculate the fold change using pP-TFBS **(C)** or TFBS-D1 **(D)** as respective controls. For **(B)**-**(D)**, student’s T-test (two tailed) was used for the pairwise comparisons of luciferase activity. P-values are reported as: *<0.05, **<0.01, ***<0.001. Mean and standard deviation are shown for each plasmid.

The enhancer activities were significantly reduced in SSc compared to control fibroblasts. We performed serial deletion using the pP-TFBS to further map the enhancer region (**Figure 3C**). A 91bp deletion (TFBS-D1) slightly increased the enhancer activity (108 ± 13%), while further deletion of 92bp (TFBS-D2) retained only 65 ± 18% of the activity. The enhancer activity was almost fully abolished when an additional 104bp (TFBS-D3) and 231bp (TFBS-D4) were deleted (6 ± 0.5%, p<0.001 for both). Deletion of the proximal 515bp (TFBS-D5) and 425bp (TFBS-D6) completely abolished enhancer activity (3 ± 0.8% and 4 ± 0.3%, p<0.001 for both), while proximal 321bp deletion (TFBS-D7) resulted in 46 ± 2%(p=0.005) enhancer activity. Lastly, deletion of proximal 193bp (TFBS-D8) resulted in stronger enhancer activity compared to the parental TFBS clone (149 ± 14%, p=0.005). This mapped the distal enhancer to an area between -584 and - 261bp from the distal TSS.

We further refined the enhancer region by serial deletion of the 515 bp TFBS-D1 clone (−584 to -68) from both 5’ and 3’ ends and tested the enhancer activity in control lung fibroblasts. Among all deletions, a 343bp element (−584 to -242, TFBS-C) resulted in a 5.8-fold (±0.6) increased activity compared to TFBS-D1 (p=<0.001). Thus, the enhancer appears to reside in this 343bp region (**Figure 3D**).

### Distal enhancer activity is partially mediated through AP-1

To identify transcription factors that may mediate the enhancer activity, we mined the UCSC genome browser and identified binding sites for multiple transcription factors (**supplemental Figure 1**). Since AP-1 is known to be an important transcription factor in extracellular matrix metabolism [23], and also has multiple known binding sites, we searched for an AP-1 binding site within our 343bp enhancer region using PROMO [32]. Two clusters of AP-1 binding sites were identified at -525 to -520 (site 1) and -358 to -353 (site 2)(**Figure 4A**). To determine if one or both of the AP-1 sites were functional, we performed site-directed mutagenesis of each site individually and tested the enhancer activity. Similar to the reduced activity for TFBS in SSc, we observed significantly lower enhancer activity in IPF fibroblasts compared with controls for the pP-TFBS-C (IPF: 110.1 ± 15.6 and control: 217.4 ± 24.4 fold, p<0.001)(**Figure 4B**). Mutation of site 1 (pP-M1) retained the enhancer activity in control fibroblast (207.0 ± 8.1 fold) and resulted in a slightly reduced activity in IPF fibroblasts (94.2 ± 11.5 fold). Mutation of site 2 (pP-M2) partially abolished the activity in both control (649 ± 4.9 fold) and IPF fibroblasts (24.0 ± 2.4 fold, p<0.001 for both). Co-expression of a *FOS* expression plasmid with the 343 bp enhancer led to 5.4 ± 0.5 fold and 4.3 ± 0.3 fold (p<0.001 for both) increase in enhancer activities for control and IPF fibroblasts. Similar increases were observed in control and IPF fibroblasts for the M1 mutation by *FOS*. There was only about two-fold increase in enhancer activity for both control and IPF fibroblasts in the M2 mutation condition (**Figure 4C**).

**Figure 4.**
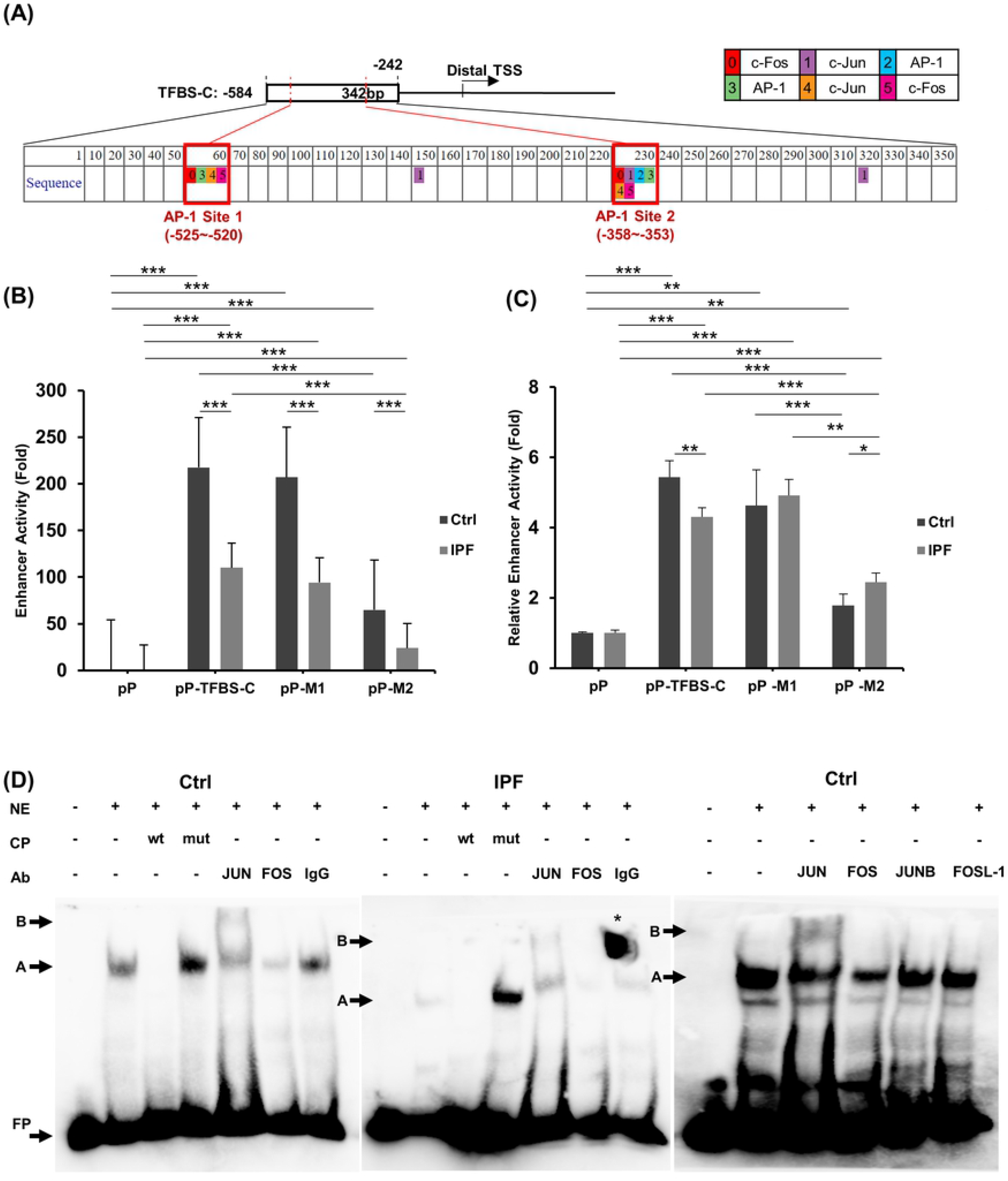
Functional AP-1 binding site associated with the distal enhancer. **(A)** Locations of AP-1 binding site clusters in the distal enhancer identified using the PROMO prediction online tool are shown. **(B)** Enhancer activities of the AP-1 site specific mutations in the distal enhancer created by site-directed mutagenesis in control (Ctrl) and idiopathic pulmonary fibrosis (IPF) lung fibroblasts are shown. Enhancer activity was calculated using pGL2promoter (pP) as a control. **(C)** Increases of enhancer activity by over-expressing FOS in control and IPF fibroblasts is shown. A FOS expression plasmid driven by a CMV promoter was co-transfected into fibroblasts with each enhancer reporter plasmid using control and IPF fibroblasts. Co-transfection with pcDNA3 was used as a vector control and for calculating the fold increase in enhancer activity by FOS. Transfections were performed in quadruplicates for both **(B)** and **(C)** and student’s T-test (two tailed) was used for pairwise comparisons of luciferase activity. P-values are reported as: *<0.05, **<0.01, ***<0.001. Mean and standard deviation are shown for each plasmid. **(D)** Binding of JUN and FOS to the AP-1 site 2 analyzed using Electrophoretic Mobility Shift Assays (EMSA) and supershift analysis is shown. Nuclear proteins isolated from control (left and right gels) and IPF (middle gel) fibroblasts were used for the binding assay with a biotin-labeled probe spanning the AP-1 site 2. The addition of unlabeled wildtype (wt) or mutated (mut) probe, and specific antibodies for IgG were used for the supershift analysis and are labeled. The band corresponding to the specific complex, supershifted complex by JUN, and free labeled probe are marked as A, B, and FP, respectively. The signal between lanes 6 and 7 denoted with a * is presumed to be an artifact as it is not situated within one lane.

We further tested the direct binding of AP-1 factors to the enhancer using a labeled probe spanning the functional site 2 of AP-1 and nuclear proteins isolated from control (**Figure 4D** left and right gel) and IPF (**Figure 4D** middle gel) lung fibroblasts. As shown in (**Figure 4D**, left and middle gel, lane 2), addition of the nuclear proteins resulted in a complex formation (labeled as A) compared to probe alone (FP). Addition of 50-fold unlabeled wildtype probe fully out-competed the labeled complex (**Figure 4D**, left and middle gel, lane 3) while addition of unlabeled probe with a mutated AP-1 site did not abolish the complex A formation. Instead, there was an increase of the complex A intensity with this condition (**Figure 4D**, left and middle gel, lane 4). Supershift experiments with an antibody specific for JUN resulted in a higher molecular weight complex (**Figure 4D**, left and middle gel, lane 5, right gel, lane 3 label B), while use of an antibody specific for FOS reduced the intensity of complex A, indicating an additional complex formation between complex A and FOS antibody (**Figure 4D**, left and middle gel, lane 6, right gel, lane 4). In the control, rabbit IgG did not reduce the complex A intensity (**Figure 4D**, left and middle gel, lane 7). Nuclear proteins isolated from IPF fibroblasts (**Figure 4D**, middle gel) resulted in similar pattern complex formations as that isolated from control fibroblasts (**Figure 4D**, left gel) at a much lower intensity. Analysis with additional AP-1 TF including JUNB and FOSL1 showed no additional supershifted complex and very little changes in the complex A intensity (**Figure 4D**, right gel, lanes 5 and 6). This suggests that JUN and FOS directly bind to the AP-1 site located at the -358 to -353 position in the distal enhancer.

### Reduced expression of JUN and FOS in IPF lungs and direct correlations to RXFP1 gene expression

Microarray whole lung tissue gene expression was performed for 108 controls and 160 IPF in the LGRC dataset [29] for *FOS* and *RXFP1*. The demographic and clinical characteristics are shown in **Supplemental Table 2**. Patients with IPF had lower expressions of *FOS* compared to controls (15.2 ± 1.7 and 13.5 ± 1.6 normalized hybridization signal for controls and IPF, p<0.001)(**Figure 5A**). The expression levels of *FOS* were positively correlated with *RXFP1* in patients with IPF (R^2^ =0.060, p=0.002)(**Figure 5B**). The expression levels of *JUN* analyzed using RNAseq from 22 controls and 22 patients with IPF showed reduced levels in IPF compared to controls (1377 ± 39.5 and 1053 ± 535 fPKM for controls and IPF, p=0.001)(**Figure 5C**). The expression levels of *JUN* were also correlated to the levels of *RXFP1* analyzed using RNAseq (R^2^ =0.365, p<0.001) **Figure 5D**) in IPF.

**Figure 5.**
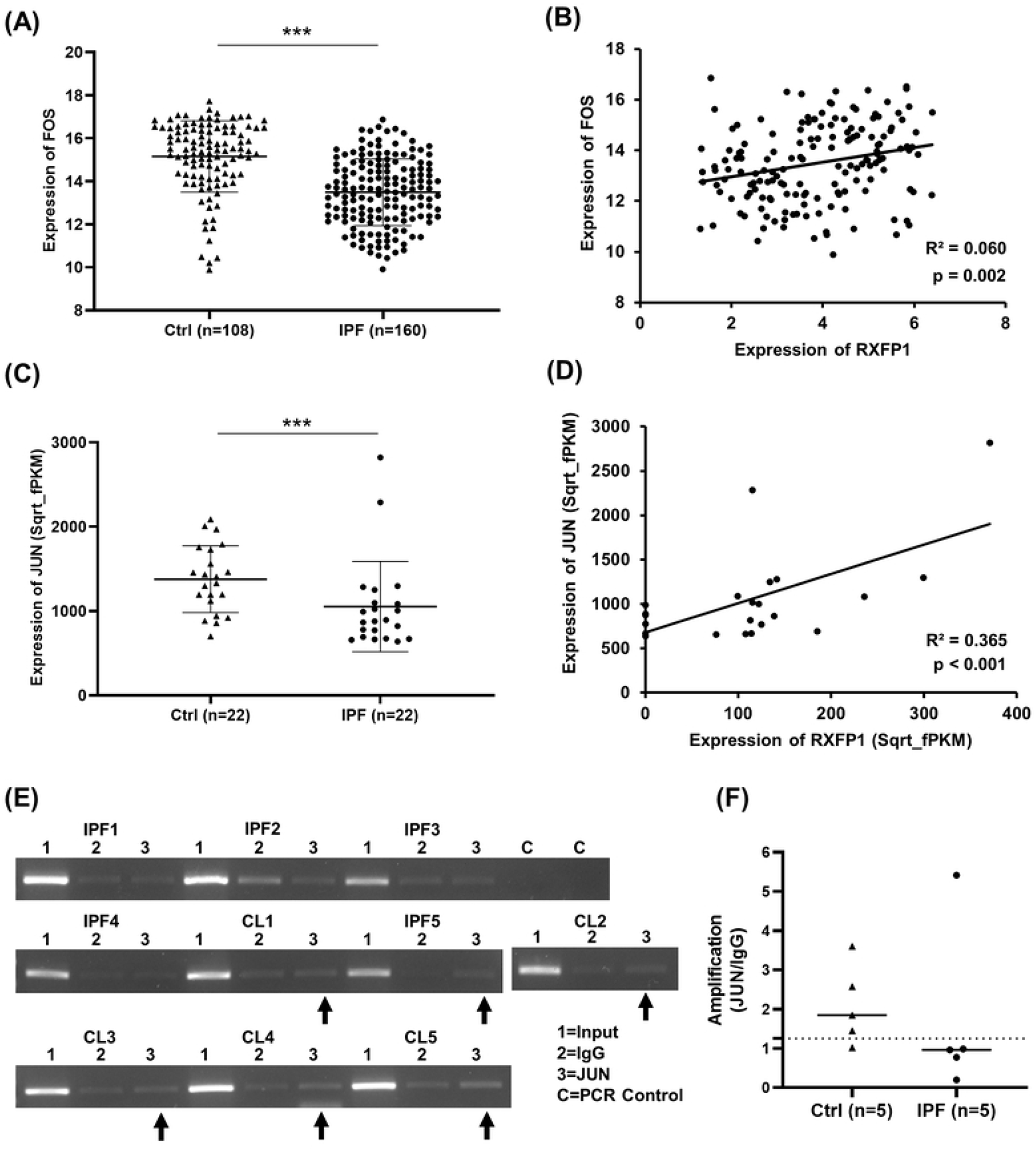
Lower expression of *JUN* and *FOS* and positive correlation to RXFP1 in control and IPF lungs. Lung tissue expression levels of *FOS* analyzed using microarray **(A)** and *JUN* analyzed using bulk RNA sequencing **(C)** from the publicly available Lung Genomics Research Consortium (LGRC) gene expression dataset (GEO accession GSE47460; http://www.lung-genomics.org/) were compared between control (108 subjects for *FOS* and 22 subjects for *JUN*) and idiopathic pulmonary fibrosis (IPF) (160 subjects for *FOS* and 22 subjects for *JUN*). The mean and standard deviation for each group and Mann-Whitney U test p-values are shown. Correlation of *FOS* **(B)** and *JUN* **(D)** gene expression levels with *RXFP1* was analyzed in IPF lungs (160 subjects for *FOS* and 22 subjects for *JUN*) using linear regression and the R^2^ and p-value are shown. **(E)** Chromatin Immunoprecipitation (ChIP) analysis of 5 pairs independent control and IPF lung fibroblasts for JUN binding to the *RXFP1* distal enhancer is shown. The distal enhancer region from -394 to -245 of distal TSS was amplified using pulldown with DNA samples. Input and IgG pulldown products were used as controls. **(F)** Densitometry analysis of the PCR amplification from **(E)** using ImageJ is shown. Positive binding by JUN to the distal enhancer was calculated by JUN/IgG density using an arbitrary cutoff of 1.25 (25% increase in binding for JUN). Student’s T-test (two tailed) was used for the pairwise comparison between IgG and JUN, and significance was not met.

ChIP analysis was performed using an antibody specific for *JUN* and 10 independent lung fibroblast lines (5 control and 5 IPF). Densitometry analysis of the PCR amplification products with primers spanning the AP-1 binding site 2 showed that 4/5 of the control lines had pulldown signal using an arbitrary cutoff of a 25% increase in PCR amplification for the pulldown ratio (JUN/IgG) >1.25, while only 1/5 IPF lines was positive (**Figure 5E**). The two-group comparison did not meet statistical significance (1.67 ± 2.11 vs 2.10 ±1.02)(**Figure 5F**). These findings suggest that in comparison to control fibroblasts, there may be lower JUN binding to the functional AP-1 site of the *RXFP1* distal regulatory region in IPF fibroblasts.

## Discussion

We have identified a strong enhancer within the distal regulatory region of *RXFP1*, which had reduced activities upon introduction into fibrotic lung fibroblasts compared to controls. The enhancer activity was partially mediated by AP-1, with lower expression of *JUN* and *FOS* in lungs from patients with IPF compared to controls and lower binding of JUN to the enhancer region in IPF fibroblasts. Furthermore, expression levels of *JUN* and *FOS* were positively correlated with *RXFP1* expression in lung tissue from patients with IPF. This is the first study to systemically analyze the regulatory elements of *RXFP1*, thus providing molecular insights into transcriptional regulation of this important protein in lung fibroblasts.

A number of studies support relaxin as a potent anti-fibrotic agent [7-11, 33, 34]. Relaxin enhances the degradation of ECM in tissues by upregulating members of the matrix metalloproteinase (MMP) family [35]. The failed clinical studies for relaxin-based treatments in SSc patients [12] may be related to reduced expression of *RXFP1* in fibroblasts of these patients, which would abrogate their responsiveness to relaxin [17-19, 31]. Patients with IPF and SSc with higher *RXFP1* expression in their lungs have better pulmonary function, supporting the pathophysiologic relevance of this locus in fILDs [17]. *In vitro* silencing of *RXFP1* results in insensitivity to exogenous relaxin, an effect which is reversed by enhancement of *RXFP1* expression in both control and IPF lung fibroblasts [17]. In this context, upregulation of *RXFP1* may serve as a therapeutic option that would help to restore the responsiveness to relaxin-based therapies in fibrotic tissues [36]. Our study suggests that transcriptional modulation of *RXFP1* in fibroblasts from patients with fILD may be one of the strategies to restore *RXFP1* expression and the responsiveness to relaxin-based antifibrotic therapies in patients with IPF and SSc.

AP-1 is ubiquitously expressed in different cells and tissues and plays important roles in multiple cellular processes including proliferation, differentiation, senescence, and cell death [21, 37]. The AP-1 superfamily consists of four subfamilies, including FOS, JUN, ATF, and MAF, which exert their functions as homo- or hetero-dimers formed through their basic leucine-zipper (bZIP) motifs. The dimers formed with different AP-1 proteins are often associated with differential transcriptional regulation of target genes [38]. In general, the dimer of FOS and JUN is associated with positive gene regulation, while other family members such as JUNB act as negative transcriptional regulators [38]. Context dependent regulation by AP-1 transcription factors is also reported [37, 39]. AP-1 transcription factors can also preferentially bind to distal enhancers instead of promoters in regulating target genes [40], supporting the finding from this study. We identified FOS and JUN as positive regulators for the *RXFP1* gene distal enhancer in lung fibroblasts. We also found that their expression levels were reduced in IPF lungs. By upregulating these transcription factors in IPF fibroblasts we may be able to restore *RXFP1* expression and thus responsiveness to relaxin-based therapeutics in fibrotic fibroblasts.

Conversely, FOSL2, a member of the AP-1 FOS subfamily has been shown to exert profibrotic effects. Transgenic *Fosl2* mice develop spontaneous lung fibrosis with *Fosl2*-expressing macrophages promoting lung fibrosis [41, 42]. Interestingly, in the LGRC expression dataset, expression levels of *FOSL2* and *RXFP1* were negatively correlated (data not shown). Therefore, the differential effects on lung fibrosis between JUN and FOS from this study in fibroblasts and the *FOSL2* expression in mice macrophages illustrates the complexity of AP-1 family functions in lung fibrosis. Additionally, we found that miR-144-3p downregulates *RXFP1* expression through 3’-untranslated region and JUN was required for constitutive miR-144-3p expression in lung fibroblasts, suggesting distinct function may be associated with the same AP-1 factor depending on their partners for dimerization. Although it is out of the scope of this study, systematic analysis of different AP-1 members in regulating, positively or negatively, *RXFP1* expression is important for understanding the transcriptional regulation of this gene. The lack of regulatory functions in the proximal region is surprising, and may be due to the distal enhancer regulating the proximal region over a long range, for example through chromatin conformation changes [43]. As reviewed by Bejjani and colleagues, genome-wide analysis has shown that AP-1 commonly binds the distal enhancers and regulates distant genes [40]. Analysis of the chromatin architecture in the *RXFP1* locus will be essential to determining whether AP-1 mediates distant control of the weak proximal regulatory region of *RXFP1* through this mechanism. In addition, reduced AP-1 binding to the *RXFP1* enhancer in IPF fibroblasts maybe due to the masking of AP-1 binding site by differential DNA methylation in this locus in IPF fibroblasts. Therefore, characterization of the epigenetic changes in fibrotic fibroblasts are warranted.

Our study does have some limitations. First, the analysis of *RXFP1* regulatory elements was mainly performed using primary lung fibroblasts from control, IPF and SSc lungs. We showed reduced direct binding of JUN to the *RXFP1* enhancer in lung fibroblasts using ChIP assay and positive correlation of *JUN* and *FOS* expression with *RXFP1* in IPF whole lung expression. However, the expression in whole lung may mask the cell-type specific expression differences of these genes. Second, we analyzed the *in vivo* binding of AP-1 to the *RXFP1* enhancer using 5 pairs of control and IPF lung fibroblast lines. Since the fibroblasts isolated from lungs are often heterogeneous [44], analysis in additional independent fibrotic and control fibroblast lines is warranted. Third, the AP-1 family consists of a large number of different transcription factors with some distinct and similar functions [21, 37]. We only focused our analysis on JUN and FOS. Comprehensive analysis of other AP-1 family members in fibroblast *RXFP1* regulation is important.

In conclusion, we identified a distal enhancer of *RXFP1* with differential activity in fibrotic lung fibroblasts involving AP-1 transcription factors. Our study provides insight into the reduced expression of *RXFP1* in patients with IPF and may support efforts to restore the effectiveness of relaxin-based therapeutics in fILD through the upregulation of *RXFP1* transcription.

## Acknowledgements

Funded by National Institute of Arthritis and Musculoskeletal and Skin Diseases (NIAMS) R21AR076024-01 (YZ) and pilot grant from 5P50AR060780-08 (YZ); Ministry of Science and Technology in Taiwan (MOST) 108IPFA0900005 (TYC). The funders had no role in study design, data collection and analysis, decision to publish, or preparation of the manuscript.

## Authors’ contribution

Y.Z., and T.Y.C.: study conceptualization; Y.Z., T.Y.C., and X.L.: experimental planning; T.Y.C., X.L. and K.H.: carried out the experiments; Y.Z., T.Y.C., X.L. and G.C.G.: data analysis; Y.Z., T.Y.C., X.L., G.C.G., C.H.H., J.T., H.B. and D.J.K.: critical interpretation of experimental data; Y.Z., T.Y.C., and X.L.: manuscript drafting; Y.Z. T.Y.C., and G.C.G: manuscript revision and editing; All authors provided critical review of the manuscript.

